# Building a high-resolution in vivo minimum deformation average model of the human hippocampus

**DOI:** 10.1101/160176

**Authors:** Nina Jacobsen, Julie Broni Munk, Maciej Plocharski, Lasse Riis Østergaard, Lars Marstaller, David Reutens, Markus Barth, Andrew L. Janke, Steffen Bollmann

## Abstract

**Objective:** Minimum deformation averaging (MDA) procedures exploit the information contained in inter-individual variations to generate high-resolution, high-contrast models through iterative model building. However, MDA models built from different image contrasts reside in disparate spaces and their complementary information cannot be utilized easily. The aim of this work was to develop an algorithm for the non-linear alignment of two MDA models with different contrasts to create a high-resolution in vivo model of the human hippocampus with a spatial resolution of 300 μm.

**Methods:** A Turbo Spin Echo MDA model covering the hippocampus was contrast matched to a whole-brain MP2RAGE MDA model and aligned using cross-correlation and non-linear transformation. The contrast matching algorithm followed a global voxel location-based approach to estimate the relationship between intensity values of the two models. The performance of the algorithm was evaluated by comparing it to a non-linear registration obtained using mutual information without contrast matching. The complimentary information from both contrasts was then utilized in an automated hippocampal subfield segmentation pipeline.

**Results:** The contrast of the Turbo Spin Echo MDA model could successfully be matched to the MP2RAGE MDA model. Registration using cross correlation provided more accurate alignment of the models compared to a mutual information based approach. The segmentation using ASHS resulted in hippocampal subfield delineations that resembled the tissue boundaries observed in the Turbo Spin Echo MDA model.

**Conclusion:** The developed contrast matching algorithm facilitated the creation of a high-resolution multi-modal in vivo MDA model of the human hippocampus. This model can be used to improve algorithms for hippocampal subfield segmentation and could potentially support the early detection of neurodegenerative diseases.

## Introduction

The hippocampus is a key structure involved in long-term memory retrieval, spatial processing, and associative learning (Burgess et al., 2002; Hartley et al., 2014; O’Keefe and Nadel, 1978; Scoville and Milner, 1957). Particularly, hippocampal volume decrease correlates with memory loss in neurodegenerative diseases such as Alzheimer’s disease (Weiner et al., 2013). Recent studies suggest that specific substructures of the hippocampus, the subfields or laminae, are differentially affected during disease progression and that volumetric measures of individual subfields are more sensitive markers of disease (Boutet et al., 2014; Henry et al., 2011; Kerchner et al., 2010; La Joie et al., 2013; Maruszak and Thuret, 2014; Pluta et al., 2012). In particular, detailed volumetric segmentation of the hippocampus could improve diagnosis and prognosis of neurodegenerative diseases (Suppa et al., 2016, 2015), or hippocampal sclerosis in patients with mesial temporal lobe epilepsy (Coras et al., 2014). Due to the convolved anatomy of these substructures, a high spatial resolution and a high signal-to-noise ratio is necessary in single subjects to obtain an accurate segmentation. This is challenging with standard scanning and analysis protocols. Therefore, automatic segmentation algorithms of anatomical structures, such as the hippocampus, generally use atlas-based approaches (Cabezas et al., 2011; González-Villà et al., 2016).

An atlas is the combination of a template image that shows structures of interest, and segmented labels. The details that can be differentiated in the template image determine final segmentation results; therefore, high-resolution data from post-mortem samples (Iglesias et al., 2015; Wisse et al., 2016a; Yushkevich et al., 2009) or high-resolution in vivo datasets (Goubran et al., 2014; Yushkevich et al., 2015) have been used to construct hippocampus atlases. However, the resolution achievable in single subjects in vivo is constrained by scan time and motion artefacts. Ex vivo data can be acquired at higher resolution but the applicability of the resulting atlases is limited due to geometric distortions, shrinkage and a limited representation of younger populations due to the inclusion of elderly subjects with hippocampal atrophy (Iglesias et al., 2015). In this work we aim to overcome these problems by constructing a high-resolution multi-modal in vivo model of the human hippocampus using minimum deformation averaging (MDA).

Minimum deformation averaging techniques combine data from a group of representative participants to form a high-resolution synthetic model, which captures the average morphology of the population used in the model generation (Grabner et al., 2006; Janke and Ullmann, 2015). The average is generated by repeatedly aligning single-subject data using non-linear registrations. Due to the averaging procedure, an MDA model will only contain structures that are anatomically consistent and can be aligned successfully across the population. Consequently, such a model enhances resolution, contrast- and signal-to-noise ratios compared to single-subject data. The feasibility of using MDA in an automatic atlas-based segmentation has already been demonstrated for the entire structure of the hippocampus (Grabner et al., 2006).

For the automatic segmentation of hippocampal subfields it is crucial to use both T1- (T1w) and T2- weighted (T2w) images, as these contrasts contribute complementary structural information (Winterburn et al., 2013). T1w images allow one to delineate the grey matter of the hippocampus from the white matter of the surrounding temporal lobe, whereas T2w images show enhanced contrast between the individual hippocampal layers (Winterburn et al., 2013). A high-resolution multi-modal in vivo model of the human hippocampus could therefore be obtained by combining T1w and T2w MDA models, generated as described in (Janke et al., 2016). However, due to the modality-specific averaging procedure, these models are in disparate anatomical spaces and require non-linear alignment before they can be fused into a single multi-modal model.

Transformation parameters to align multi-modal images are commonly estimated using mutual information as the similarity measure. Mutual information describes the statistical dependence between image intensities (Maes et al., 1997) and is a robust option for rigid multi-modal registration (Lu et al., 2008). However, the use of mutual information in non-rigid registration scenarios can cause unwanted deformations and misalignment due to a loss of statistical consistency at smaller scales (Andronache et al., 2008). For non-rigid registration the cross-correlation coefficient is a computationally efficient and more robust option (Andronache et al., 2008). However, the cross-correlation coefficient requires equal contrast in both images, which renders it impracticable for multi-modal applications. To overcome this limitation, Andronache and colleagues (2008) proposed a method where different modalities are mapped into a common contrast space. The result of this is the creation of a pseudo-modality, which only shows image features common in both images. Details only visible in one of the images are consequently ignored, making this method not ideal for our purpose.

Here, we propose the application of a contrast matching algorithm to estimate non-linear registration between images of different modalities using the cross-correlation coefficient. Therein, the contrast of one image is transformed into the contrast of an image of a different modality. This then allows the application of the cross-correlation coefficient to estimate a non-linear registration for high-resolution data. Importantly, features present in only one image are retained and can be later combined to formulate a multi-modal MDA model. To demonstrate the feasibility of this method, we perform a combined segmentation of the hippocampal subfields using a Turbo Spin Echo (TSE) and an MP2RAGE MDA model.

## Methods

### Data acquisition

Imaging data were acquired at the Centre for Advanced Imaging at the University of Queensland using a 7T whole body MRI scanner (Siemens Healthcare, Erlangen, Germany) equipped with a 32-channel head coil (Nova Medical, Wilmington, MA, USA) and an SC72 gradient system providing a maximum gradient strength of 70 mT/m and a slew rate of 200 mT/m/s. To improve the B_0_ field homogeneity 3^rd^order shimming was employed.

Whole-brain T1w images were acquired using a prototype MP2RAGE sequence (Siemens WIP 900) in 48 participants (16 female, average age 31.1±8.6) with a range of isotropic resolutions: 0.5 mm (18 participants), 0.75 mm (21 participants), 1.0 mm (8 participants) and 1.25 mm (1 participant). Common image parameters were TR = 4330 ms, TI1/TI2 = 750/2370 ms, TE = 2.8 ms, flip angles = 5 and 6 degrees, and GRAPPA = 3.

T2w images were acquired using a 2D TSE sequence covering a slab orthogonally to the main axis of the hippocampus. 26 individuals (13 female, average age 26.8±3.9) were imaged using 3 repetitions and the following parameters: resolution = 0.4×0.4×0.8 mm, flip angle = 134 degrees, and TR = 10.3s. Before the atlas creation, the three TSE acquisitions per participant were averaged into one dataset to increase the signal-to-noise ratio. Then the TSE dataset was resampled to an isotropic voxel size of 0.4 mm.

### Model generation

Probabilistic models were created for each image contrast individually using MDA as described in (Janke et al., 2016). The MP2RAGE (Marques et al., 2010) denoised images (O’Brien et al., 2014b, 2014a) were used in a fitting strategy consisting of one linear fit to the evolving internal model followed by a hierarchical series of non-linear grid transforms. These transforms started with a step size of 32 mm followed by 16 mm, 12 mm, 8 mm, 6 mm, 4 mm, 2 mm, 1.5 mm, 1 mm, and finished with 0.8 mm. The fitting steps used progressively de-blurred data with a 3D kernel full width at half maximum of half the current step size. Twenty iterations at the first 5 fitting stages, 10 iterations at the next 3 stages and 5 iterations at the last 2 stages were performed using the ANIMAL algorithm (Collins et al., 1994). As the step size decreased, the resolution of the evolving model was increased, starting with a step size of 1.0 mm and finishing with 0.3 mm. It is possible to increase the resolution to this point as voxels from different participants contain overlapping information, which allows the extraction of sub-voxel boundary information. A robust averaging process, where outliers beyond a standard deviation are down-weighted, was used throughout to create the MP2RAGE MDA model (Bollmann et al., 2017a)(Figure 1, right).

**Figure 1.**
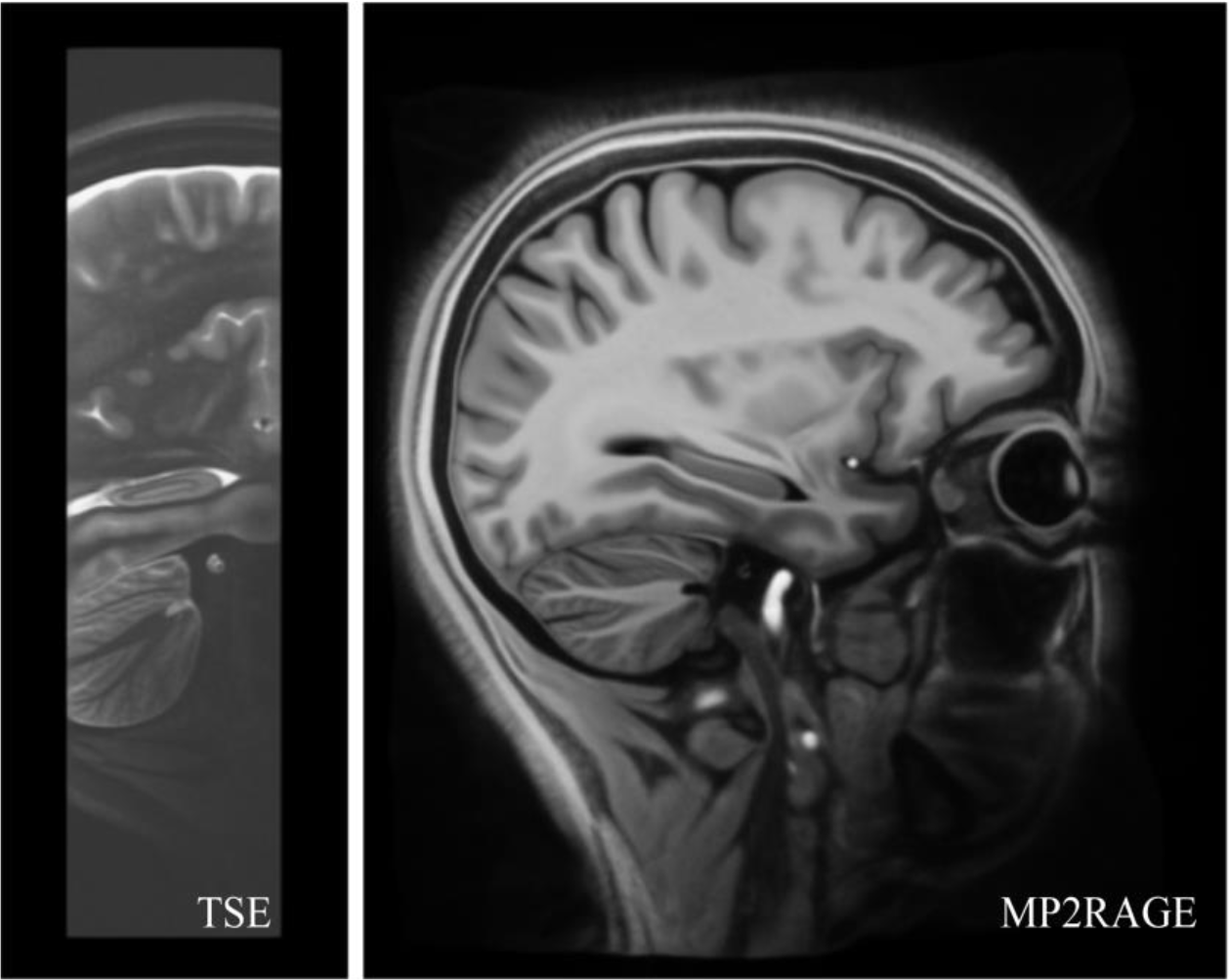
Sagittal view of the TSE (left) and the MP2RAGE MDA model (right). Both models show improved contrast- and signal-to-noise ratios compared to single-subject data with a high amount of detail in the hippocampus.

The averaged TSE data were first normalized to an intensity range between 0 and 100 by a histogram critical threshold clamping. Then, the intensity was clamped between 0 and 30 to increase the contrast further. The hierarchical series of non-linear grid transforms used the same step sizes as for the MP2RAGE model but finished with a step size of 0.4 mm. The final resolution of the TSE MDA model was 0.2 mm (Bollmann et al., 2017b) (Figure 1, left).

### Contrast matching algorithm

The aim of the contrast matching algorithm (CMA) was to estimate a non-linear transfer function of intensity values from one image modality to the other. The CMA determines the intensity relationship between the two contrasts and then uses this relationship to match them. Prior to estimating the transfer function, the spatial alignment between the two models needs to be maximized. Therefore, the TSE model and the MP2RAGE model were first manually aligned followed by a rigid-body registration using mutual information as the similarity measure (Figure 2A). Then both models were normalized to an intensity range between 0 and 100 by histogram clamping. Next, FSL bet (Smith, 2002) was used to create a whole-brain mask on the basis of the MP2RAGE model (Figure 2B). Last, the MP2RAGE model was smoothed using a uniform filter (Oliphant, 2007) with a kernel size of 21 voxels (6.3 mm) to reduce the influence of noise and remaining misalignment between the two contrasts (Figure 2B).

**Figure 2.**
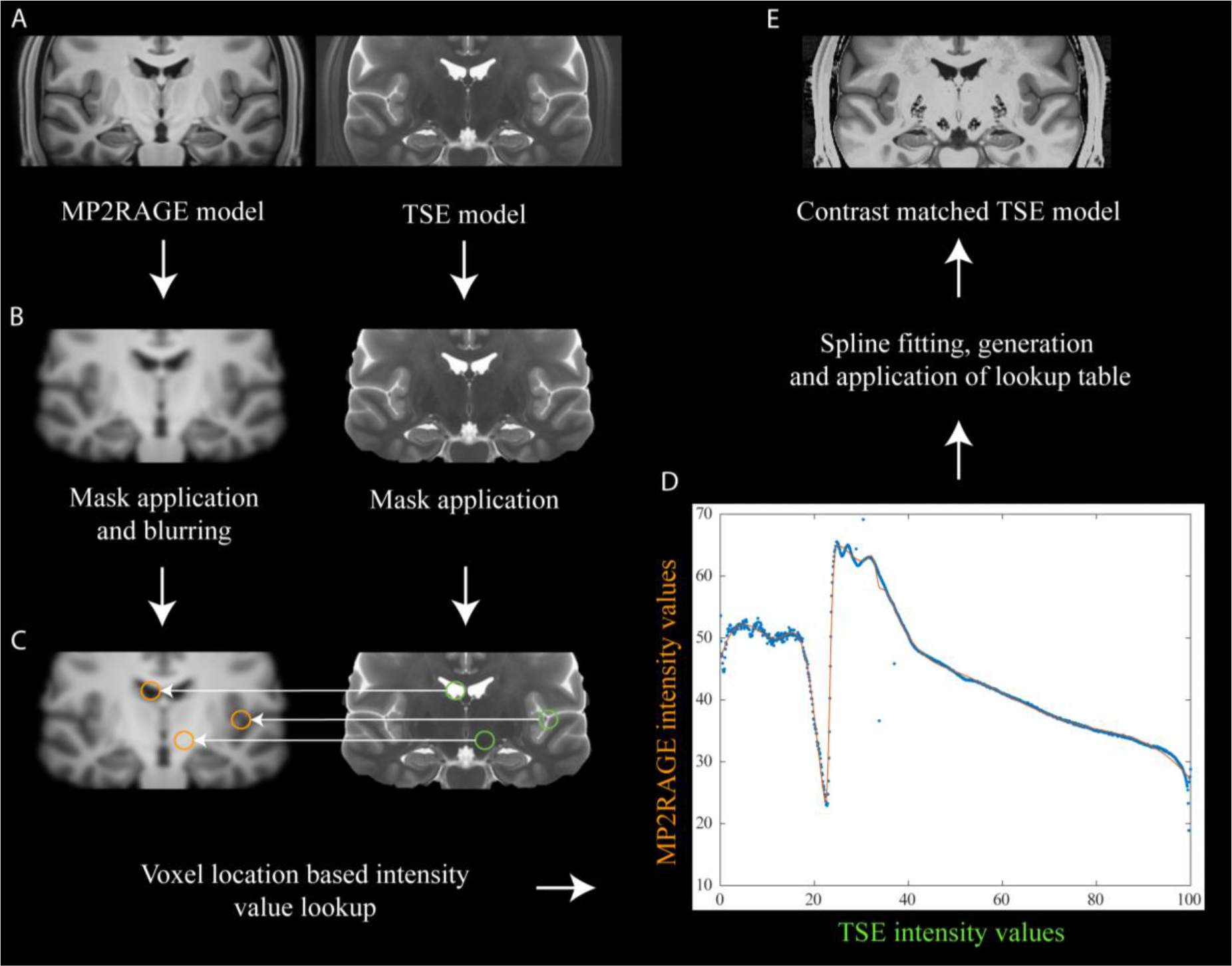
Schematic illustration of the contrast matching algorithm. The algorithm takes two modalities as input and returns one contrast-matched to the other. Both inputs are normalized (A) and masked (B), and the target contrast is blurred (B). The voxel intensity relationship is determined on the basis of voxel location (C), and a spline fit is used to yield a continuous transfer function (D, estimates: blue, fit: red). From this function a lookup table is generated which is applied to transform the TSE model’s contrast to an MP2RAGE contrast (E).

The intensity transfer function was determined by identifying the most frequent intensity value in the MP2RAGE model for every unique intensity value in the TSE model (Figure 2C). In case of multiple intensity values with highest frequency, an average of these was computed to yield the matching intensity value. A univariate cubic spline (Oliphant, 2007) with a smoothing factor of 1200 was then fit to the voxel intensity relationship to estimate a continuous transfer function (Figure 2D). Using this continuous transfer function, a lookup table consisting of the intensity values of the TSE contrast and the matching intensity values of the MP2RAGE contrast was generated. Finally, the TSE model’s contrast was transformed to the MP2RAGE contrast utilising this lookup table (Figure 2E).

The CMA is available on GitHub (https://github.com/NIF-au/cma) and the models can be viewed on http://www.tissuestack.org or downloaded from http://www.imaging.org.au/Human7T/ (Bollmann et al., 2017a, 2017b).

### Non-linear model registration

To align the contrast-matched TSE model to the MP2RAGE model a non-linear registration using cross correlation as similarity measure was performed. For comparison, a non-linear registration without contrast matching using mutual information as the similarity measure was also carried out. The non-linear registrations had four resolution levels with 100, 70, 50 and 20 iterations, respectively. For each resolution the down-sampling rate was reduced by half, starting with a value of 8. The registrations were performed using the SyN algorithm (Avants et al., 2008). The registration quality was visually assessed in the hippocampus by overlaying with a wireframe, and quantified using a Boolean AND operator on the detected edges and computing the overlap in the hippocampus using the dice coefficient.

### Hippocampus subfield segmentation

The hippocampal subfields of the multi-modal MDA model were segmented using the Automatic Segmentation of Hippocampal Subfields (ASHS) software pipeline (Yushkevich et al., 2015) in the most recent implementation ‘fastashs-20170223’ utilising the ‘upennpmnc_20161128 atlas’ with the default settings.

## >Results

### Contrast matching

The voxel intensity relationship between the TSE and MP2RAGE contrasts was estimated using the developed CMA pipeline to match the TSE model to the MP2RAGE contrast (Figure 2). The obtained transfer function reveals a non-linear relationship featuring, for example, a prominent dip at an intensity level of about 20 in the TSE image (Figure 2D). Furthermore, a broad range of TSE intensity values (0-100) are mapped to a narrow MP2RAGE intensity value range (approximately 22-65, zeroes are excluded due to the applied background masking). The fitted spline captures the overall shape of the point distribution, but removes some variations in the TSE intensity ranges between 0-20 and 25-30. Figure 3 illustrates the two input modalities, i.e. the MP2RAGE MDA model (Figure 3A) and the TSE MDA model (Figure 3B), and the contrast matched TSE MDA model (Figure 3C). In many cases, a simple contrast inversion was performed, for example in the cerebrospinal fluid, which appears bright in the TSE model, but dark in the contrast-matched model.

**Figure 3.**
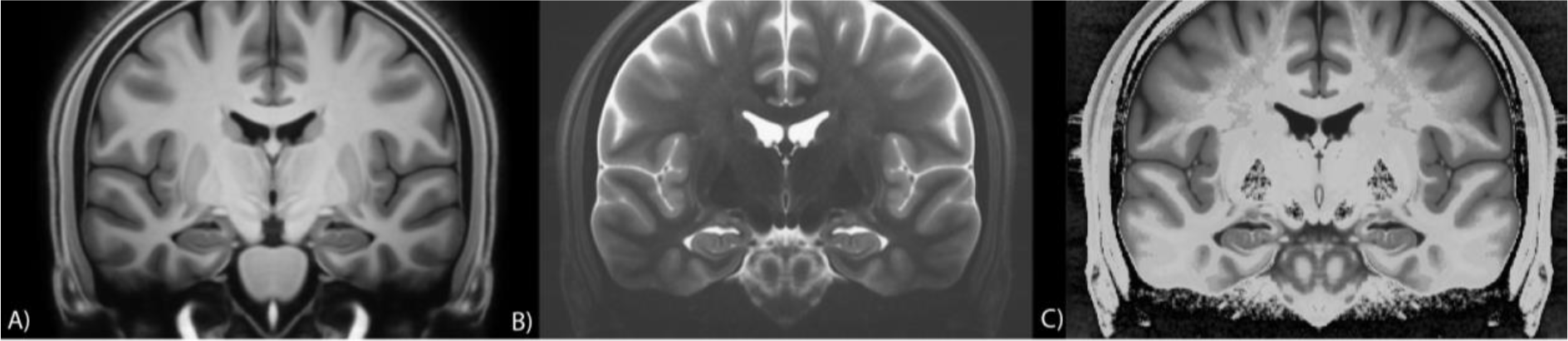
Coronal view of the MP2RAGE model (A), the TSE model (B) and the contrast matched TSE model (C).

### Linear and non-linear multi-modal registration

Figure 4 shows coronal, sagittal and transverse views of the alignment between the MP2RAGE model and the TSE model of the right hippocampus after each of the three registrations: Linear registration using mutual information, linear registration followed by non-linear registration using mutual information, and linear registration followed by non-linear registration using cross-correlation. Comparing the alignment of the MP2RAGE wireframe on each of the three registration results, the best overlap is observed with the registration result obtained by non-linear registration using crosscorrelation and CMA (Figure 4, 4^th^ column). In the coronal view (Figure 4, top row), the improved alignment is especially visible in the subiculum, located in the inferior part of hippocampus, and the inferior temporal gyrus (orange arrows). In the sagittal view (Figure 4, centre row), improved alignment is especially notable at the lateral ventricle (blue arrows). Improved alignment is similarly discernible in the transverse view (Figure 4, bottom row) where an increased overlap in the lateral ventricle is observed (green arrows). The quantitative analysis shows increasing dice scores from 0.23 for the linear registration to 0.28 for the non-linear registration using mutual information only. The highest dice score of 0.52 was obtained from the non-linear registration using CMA and crosscorrelation (Figure 5).

**Figure 4.**
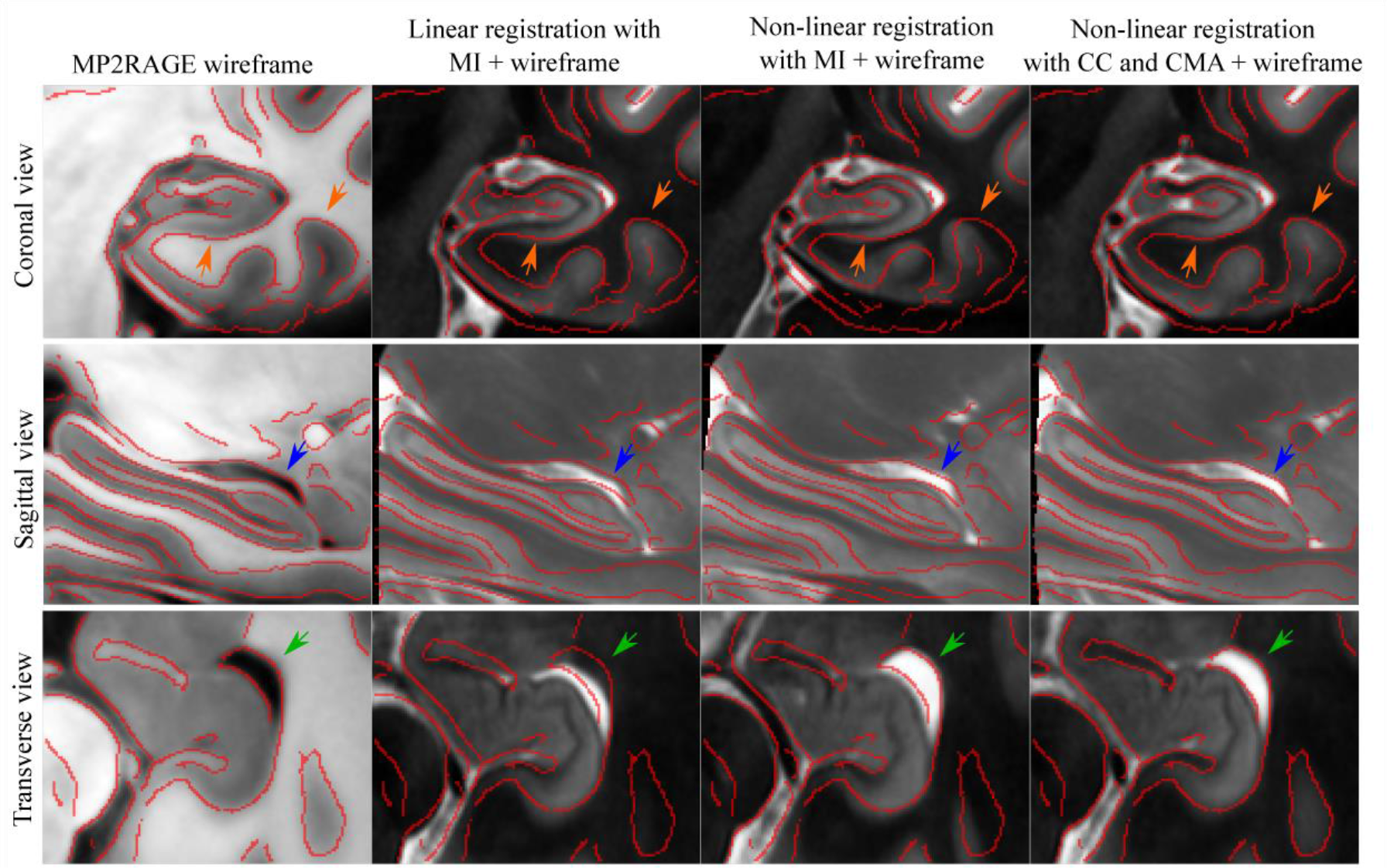
Coronal, sagittal and transversal views of the alignment in the hippocampus between the MP2RAGE model and the TSE model after three different registration pipelines. All views capture the right hippocampus. Edges in the MP2RAGE model have been detected and superimposed onto all registration results as red lines (wireframe). Differences between the registrations are highlighted by arrows: The subiculum and the inferior temporal gyrus are highlighted in the coronal view of the MP2RAGE model (orange arrows). In the sagittal and the transverse view of the MP2RAGE model the lateral ventricle is marked by blue arrows and green arrows respectively.

**Figure 5.**
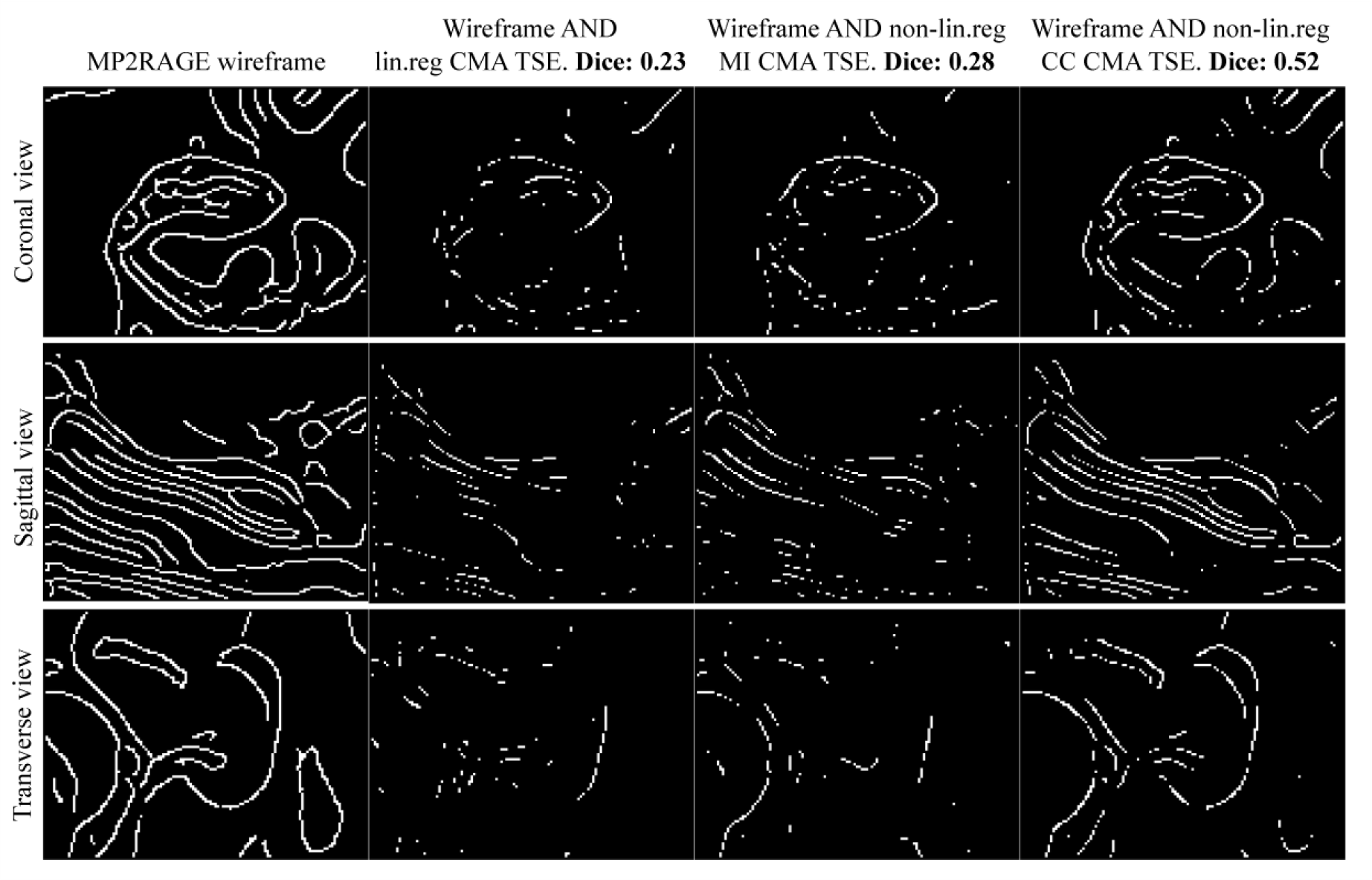
Coronal, sagittal and transversal views of the hippocampus edge alignment between the MP2RAGE model and the TSE model after three different registration pipelines. All views capture the right hippocampus. Edges in all views were detected and the Boolean AND operation was performed to show where the registered edges overlap with the MP2RAGE target edges. The overlap was quantified using the dice coefficient showing highest overlap for non-linear registration using CMA and cross-correlation.

### Hippocampus subfield segmentation

The segmentation result of the multi-modal hippocampus model utilising the ASHS pipeline is shown overlaid onto the co-registered MP2RAGE and TSE models in Figure 6. The segmented subfields illustrate the importance of using the TSE contrast in the segmentation because subfield tissue boundaries, e.g. between the dentate gyrus and the cornu ammonis field 1, are clearly visible in the TSE contrast and, hence, are segmented fully automatically using ASHS.

**Figure 6.**
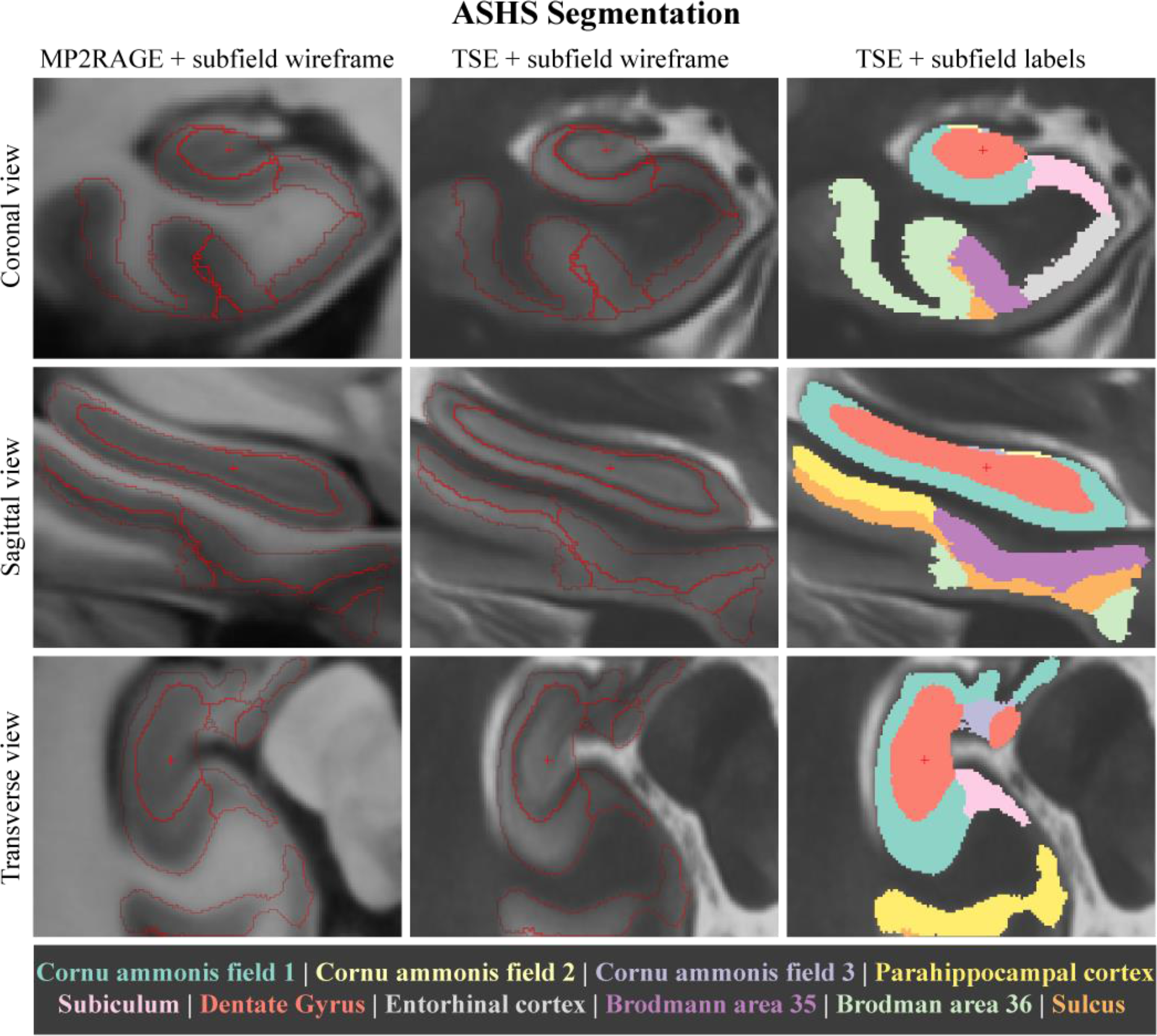
ASHS segmentation results overlaid onto the multi-modal hippocampus model. The boundary between cornu ammonis field 1 and dentate gyrus is clearly visible in the TSE model, whereas there is little contrast in the MP2RAGE model between these two subfields.

## Discussion and Conclusion

The contrast matching algorithm presented here enabled the non-linear alignment of two high-resolution MDA models using cross-correlation, outperforming mutual information as a similarity measure. The registration results are consistent with the proposal that mutual information is a suitable similarity measure when the information in the images or models is equivalent (Rueckert and Schnabel, 2014), which is not the case for the region of interest here, i.e. the hippocampus (Winterburn et al., 2013; Wisse et al., 2016b). In comparison to the algorithm by Andronache et al. (2008), details only pronounced in one of the contrasts were still preserved using the CMA. However, for some brain areas the algorithm produced contrast discrepancies. This might be caused by the estimation of a global transfer function, where a majority vote is used to ensure an unambiguous assignment of intensity values. Consequently, when different tissues in the source image have the same intensity but should map to different intensities in the target image, such as in the inferior temporal gyrus and the caudate nucleus, a mismatch in intensity values will occur. This could be overcome by estimating a local transfer function introducing a spatial dependency when estimating the voxel intensity relationship. The brain mask used in our example prevents background voxels from dominating the estimated transfer function. Furthermore, the estimated transfer function is usually not bijective, requiring a re-estimation for each direction the CMA is meant to be applied. To directly estimate a multi-modal MDA model, the CMA could be incorporated in the model building itself similar to Guimond et al. (2001). Thereby, at each averaging stage the non-linear transformations are estimated from two (or possibly more) contrasts merging information across modalities.

In conclusion, the presented algorithm (https://github.com/NIF-au/cma) enabled the building of a high-resolution multi-modal in vivo MDA model of the human hippocampus, which is available to the community at www.imaging.org.au. The obtained model allows the detailed segmentation of the hippocampal subfields utilizing complementary contrast information from T1- and T2- weighted images.

## Acknowledgements

The data acquisition was funded by the Australian Research Council Special Research Initiative: Science of Learning Research Centre [project number SR120300015]. NJ and JBM acknowledge funding from the following Danish private foundations: Otto Mønsteds Foundation, Knud Højgaards Foundation, Augustinus Foundation, Oticon Foundation, Nordea Foundation, Marie and M.R Richters Foundation, Viggo and Pedersens Foundation, and Danish Tennis Foundation. MB acknowledges funding from an Australian Research Council Future Fellowship grant [FT140100865] and DR from Discovery grant [DP140103593]. SB acknowledges funding from a UQ Postdoctoral Research Fellowship grant and an NVIDIA Hardware Seed Grant. The authors acknowledge the facilities of the National Imaging Facility at the Centre for Advanced Imaging, University of Queensland.

## Notes

**Conflict of Interest** The authors declare that they have no conflict of interest.

